# Improvement in Patient-Reported Sleep in Type 2 Diabetes and Prediabetes Participants Receiving a Continuous Care Intervention with Nutritional Ketosis

**DOI:** 10.1101/389841

**Authors:** Morgan J. Siegmann, Shaminie J Athinarayanan, Sarah J Hallberg, Amy L. McKenzie, Nasir H. Bhanpuri, Wayne W. Campbell, James P. McCarter, Stephen D. Phinney, Jeff S. Volek, Christa J. Van Dort

## Abstract

**Objective:** Sleep disruption is frequently associated with type 2 diabetes (T2D) and hyperglycemia. We recently reported the effectiveness of a continuous care intervention (CCI) emphasizing nutritional ketosis for improving HbA1c, body weight and cardiovascular risk factors in T2D patients. The present study assessed the effect of this CCI approach on sleep quality using a subjective patient-reported sleep questionnaire.

**Methods:** A non-randomized, controlled longitudinal study; 262 T2D and 116 prediabetes patientsenrolled in the CCI and 87 separately recruited T2D patients continued usual care (UC) treatment. Patients completed the Pittsburgh Sleep Quality Index (PSQI) questionnaire. A PSQI score of >5 (scale 0 to 21) was used to identify poor sleepers.

**Results:** Global sleep quality improved in the CCI T2D (p<0.001) and prediabetes (p<0.001) patients after one year of intervention. Subjective sleep quality (component 1), sleep disturbance (component 5) and daytime dysfunction (component 7), also showed improvements in the CCI T2D (p<0.01 for sleep quality and sleep disturbance; and p<0.001 for daytime dysfunction) and prediabetes patients (p<0.001 for all three components); compared to the UC T2D group after one year. The proportion of patients with poor sleep quality was significantly reduced after one year of CCI (T2D; from 68.3% at baseline to 56.5% at one year, p=0.001 and prediabetes; from 77.9% at baseline to 48.7% at one year, p<0.001).

**Conclusion:** This study demonstrates improved sleep quality as assessed by PSQI in patients with T2D and prediabetes undergoing CCI including nutritional ketosis but not in T2D patients receiving UC. The dietary intervention benefited both sleep quality and the severity of T2D symptoms suggesting that nutritional ketosis improves overall health via multiple mechanisms.

## Introduction

Sleep disruption is associated with obesity and type 2 diabetes (T2D), yet the bidirectional relationship between sleep and glucose metabolism is not fully understood. It is linked to increased diabetes prevalence in both experimental ^1-4^ and epidemiological studies ^5-7^. In addition, the severity of hyperglycemia in individuals with diabetes is associated with poor sleep quality ^8, 9, 10, 11^, short sleep duration ^8, 9, 12, 13^ and a greater tendency to develop sleep disorders including obstructive sleep apnea (OSA) ^14, 15^. Both the International Diabetes Federation (IDF) and American Diabetes Association (ADA) recommend evaluating T2D patients for sleep breathing problems especially OSA and strongly encourage treatment when found ^16, 17^.

Weight loss is one of the most effective ways to improve sleep quality, quantity [Yannakoulia, 2017 and Xanthpoulos 2018] and to treat OSA in obese patients. Lifestyle intervention induced weight loss showed significant reduction in the apnea and hypopnea indices (AHI) in conjunction with a decrease in hemoglobin A1c (HbA1c) levels in a randomized controlled trial of obese OSA patients with comorbid diabetes ^18^. Further, weight loss following bariatric surgery is effective at improving glycemic control and improving AHI in OSA patients ^19^. Intervention studies specifically targeting sleep disruption in OSA patients without any effect on weight, such as continuous positive airway pressure (CPAP) treatment, have shown contradictory results for glycemic control. Most CPAP intervention studies in T2D reported no glycemic benefit from the treatment ^20, 21^, but one study demonstrated a slight reduction in HbA1c ^22^. In contrast, CPAP studies on prediabetic OSA patients showed improvements in insulin sensitivity and glucose tolerance ^23, 24^. It is not clear from these studies whether improvement of glycemic control in conjunction with weight loss improves sleep quality or vice-versa.

A few studies have investigated the impact of dietary macronutrient composition on sleep duration and quality. Two studies reported reduction of slow wave sleep (SWS) and elevation of rapid eye movement (REM) sleep in individuals consuming higher carbohydrates (600g carbohydrate or 80% energy from carbohydrate) ^25, 26^. Another study reported the effect of a high carbohydrate (56% energy from carbohydrate) diet in reducing sleep onset latency when compared to a control diet ^27^. Studies investigating low carbohydrate diets showed the opposite effect; reduced REM ^28^, increased REM onset latency ^29^ and increased SWS ^28^, even after 4 hours of administering a very low carbohydrate meal ^28^. Collectively, these findings signify dietary carbohydrate content as an important factor in modulating sleep architecture, but extrapolation from these studies is limited since they were conducted in experimentally controlled conditions with small numbers of healthy individuals in a short time-span and with diets administered at specific time points.

Population and intervention-based studies on the overall impact of carbohydrate intake on sleep indices or sleep quality are very limited. Katagiri et al. showed reduced sleep quality in individuals consuming more carbohydrates as measured by a subjective sleep measure, the Pittsburgh Sleep Quality Index (PSQI) ^30^. Studies investigating the effect of ketogenic diet (KD) in children with sleep problems showed improvement in daytime sleepiness ^31, 32^ as well as positive changes in sleep architecture ^32, 33^. However, in one of these studies, sleep improvements were suggested to be due to weight loss rather than the KD ^33^. Despite restricted carbohydrate intake concurrent with sleep improvement in these children, SWS decreased ^33^ and REM increased ^32, 33^ which contradicts studies on carbohydrate intake and sleep architecture in adults ^25, 26, 28^. Carbohydrate restriction and ketogenic diets are widely used in the clinical management of obesity and diabetes, but studies assessing the effect of this diet on sleep are currently limited. We recently demonstrated a continuous remote care treatment for T2D including nutritional ketosis significantly improved glycemic control, weight, and cardiovascular disease risk factors and reduced diabetes medication use at one year ^34-36^.

The purpose of this study was to assess the effect of the intervention by time-interval on the global PSQI and its seven component scores as well as compared its changes with different intervention and disease categories. We also assessed the relationship between changes in the sleep parameters versus key biochemical parameters, and also investigated the correlation of pain, circadian rhythm disruption and CPAP usage versus patient-perceived sleep status. We hypothesized that the global sleep indexes would improve analogously, as improvement in other key biochemical parameters observed in the intervention.

## Materials and Methods

### Study participants and design

This study is part of a clinical trial *(Clinical trials.gov identifier: NCT02519309)* that was approved by the Franciscan Health Lafayette Institutional Review Board. Patients between age 21 and 65 years with either a diagnosis of T2D and a BMI > 25 kg/m^2^ or prediabetes and a BMI > 30 kg/m^2^ were included in this study. Detailed study design including the inclusion and exclusion criteria were previously reported ^34, 35^. Briefly, the trial was an open-label, non-randomized, controlled, longitudinal study with patients divided into three groups. The T2D and pre-diabetes patients in the continuous care intervention (CCI) regimen self-selected either on-site (CCI-onsite) or web-based (CCI-web) education delivery. Educational content and medical treatment was the same for both CCI-onsite and CCI-web. As there were no significant differences in outcomes including PSQI scores, between educational groups, they are combined for further analysis ^34, 35^. Both T2D and prediabetes CCI patients had access to a mobile health application (app) that enabled them to communicate and be continuously monitored by a team of healthcare professionals including a personal health coach and physician or nurse practitioner. Patients received individualized guidance in achieving nutritional ketosis, typically including restriction of daily dietary carbohydrates to less than 30 grams. Patients were encouraged to measure and input weight, blood glucose and blood beta-hydroxybutyrate (BHB) concentrations daily in the app. These measurements were used by the health care team for monitoring the patient’s condition (weight and glucose) and assessing carbohydrate restriction (BHB).

Separately recruited usual care (UC) T2D patients were participants in a local diabetes education program including care by their primary care physician or endocrinologist and counseling by registered dietitians; no modification to their care was made for the study. This group was observed at baseline and one year as reference for typical disease treatment and progression within the same geography and health system. (UC patients were informed that the trial had an intervention arm and could participate in that group if they chose to do so).

### Demographic and clinical variables

Patient demographic and clinical data were collected at baseline, 70 days and one year. Laboratory measures were assessed at a Clinical Laboratory Improvement (CLIA) certified laboratory. These data were initially analyzed to evaluate the safety and effectiveness of the CCI in improving diabetes status (glycemic control and medication use), weight and other metabolic factors in T2D ^34, 35^ and prediabetes patients ^36^(unpublished data, manuscript in preparation). Some of the clinical variables - weight, fasting blood glucose, HbA1c, homeostatic model assessment of insulin resistance (HOMA-IR), BHB and high sensitivity C-reactive protein (hsCRP) - were included for further analyses in this study. Usual care T2D patients were not continuously monitored for weight, blood glucose, or BHB; clinical and laboratory measures were obtained for this group only at baseline and one year.

### Pittsburgh Sleep Quality Index (PSQI)

CCI patients were administered a set of questionnaires, including the PSQI, during visits at baseline, 70 days and one year; UC participants completed questionnaires at baseline and one year. The PSQI consists of 19 validated questions assessing sleep quality and efficiency ^37^. The global PSQI score is calculated from seven component scores on subjective sleep quality (component 1), sleep latency (component 2), sleep duration (component 3), habitual sleep efficiency (component 4), sleep disturbances (component 5), use of sleep medication (component 6) and daytime dysfunction (component 7). Each question within the component is scored on a 4-point Likert scale of 0 to 3, with 3 indicating worse outcomes and the mean was calculated for each component score. The sum of the component score means generates the global PSQI score that ranges from 0 to 21. Higher global PSQI scores indicate poorer sleep. A patient with a global PSQI score ≤ 5 is considered a “good sleeper” and > 5 is categorized as a “poor sleeper” ^38^. Change in the PSQI score over time was calculated using the formula below:

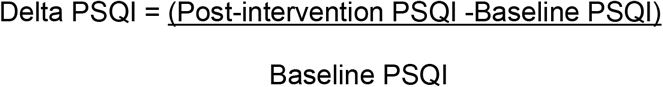

### Pain, shifted sleep chronotype and CPAP usage

Patients were classified into “pain” and “non-pain” groups based on their response to pain-related questions in both the PSQI (question 5i) and a separate questionnaire used to calculate the knee injury and osteoarthritis outcome score (KOOS). Overall KOOS results will be reported in a separate publication. Classification of patients under circadian rhythm “disrupted” and “non-disrupted” groups was based on the wake time and bedtime responses for PSQI questions 1 and 3 for compilation of component 4 (sleep efficiency). Patients were classified as having a shifted wake-up time if they reported typically waking between 11am and 2am, while those with bedtimes between 12am to 6pm were bedtime shifted. These arbitrary bedtime and wake time cut-off ranges were selected based on evening and night shift workers schedule (2nd shift - 3pm to 11pm and 3rd shift- 11pm to 7am); which causes these workers to have sleep patterns that deviate from a normal chronotype. Patients were also surveyed regarding CPAP usage and discontinuation, however detailed usage information such as CPAP pressure settings and usage compliance were not obtained making it difficult to interpret the patients OSA treatment status.

### Statistical Analyses

The questionnaires were administered by research personal and completed by patients on paper. Paper questionnaires were scanned and responses were transcribed in duplicate by an independent contract data entry firm. The patterns of missing data were assessed using Little’s MCAR test ^39^ and were found to be missing at random (MAR). Missing data were imputed by Multivariate Imputation by Chained Equations (MICE) ^40^, and Intent to treat (ITT) analyses were performed. Normality of the global PSQI and component scores was evaluated using Lilliefors test. Even after transformation, the data failed the normality test (i.e. there was a skew toward lower PSQI scores and a long tail of higher scores) (Supplemental figures 1A-C); therefore, nonparametric tests were used for analyses of PSQI scores. Results from continuous variables were expressed as mean ± standard deviation. Comparisons between groups were performed using the Kruskal-Wallis test, and comparisons within groups were performed using the Wilcoxon Sign Rank test. Tukey’s honest significant difference test was used to analyze pairwise differences among significant results from omnibus tests. McNemar’s test was used for assessing statistical significance of transitioning between ‘good’ and ‘poor’ sleeper among the CCI and UC cohorts.

Adjusted Pearson’s and Spearman correlations were calculated between changes from baseline in global PSQI and changes in metabolic-parameters. Adjusted correlations were performed while controlling for age, gender and BMI at baseline. All participants in the CCI group were stratified by sleep improvement status based on their baseline and one year global PSQI scores. Patients that were initially considered “poor sleepers” with a baseline PSQI > 5 but whose score after one year decreased to at or below the threshold of 5 were classified as *improved*. Those patients who were considered “good sleepers” at both baseline and one year were classified as *maintained*. Finally, those patients whose 1 year PSQI score was >5 (regardless of their baseline score) were classified as *not improved*. Stepwise analyses of covariance (ANCOVA) were performed between the three different CCI sleep status groups at one year with the change of the glucose-related, ketone and inflammatory markers, while controlling by age, gender and years living with diabetes. Statistical tests were performed with MATLAB R2017b using the Statistics and Machine Learning Toolbox ^41^ and the R statistical program version 3.5.0. ^42^

## Results

### Baseline participant characteristics

Details on the recruitment and extensive baseline characteristics of the CCI and UC T2D patients were previously published ^34, 35^. The demographic, glycemic, inflammatory and sleep baseline characteristics of the participants that were included for assessments of sleep are presented in Table 1. One-hundred forty-three (54.6%) CCI T2D, 61(54%) CCI prediabetes, and 53 (62.3%) UC T2D patients completed the PSQI at all expected time points. The patients who completed the trial at one year were slightly higher than those who completed the PSQI questionnaires. Some of the patients completed the study period and laboratory analysis but were unable to attend the clinic for their 70-days and one-year follow-up visits, where they are required to complete their corresponding questionnaires. The proportion of missing PSQI data were similar across the three groups with 77.61% of CCI T2D, 79.06% of CCI prediabetes and 79.24% of UC T2D completed the PSQI in all expected time points. There were no significant differences between completers and non-completers on baseline characteristics for either group at one year of the intervention (supplemental Table 1). The global PSQI and component scores did not differ significantly among the groups (CCI T2D, CCI prediabetes and UC T2D) at baseline. The proportion of participants with overall poor sleep quality was higher in the CCI prediabetes group (77.9%) compared to the CCI T2D (68.3%) and UC T2D (68.2%) groups.

**Table 1.** Baseline characteristics of participants included in the study. Baseline data were calculated using intent-to-treat (ITT

| Int Cohorts | CCI Type 2 Diabetes | CCI Prediabetes | UC Type 2 Diabe |
| --- | --- | --- | --- |
| ers, Completers, PSQI Available (n) | 262, 218, 143 | 116, 113, 61 | 87, 78, 53 |
|  | mean (S.D.) |  |  |
| years) | 53.8 (± 8.4) | 51.9 (± 9.4) | 52.7 (± 9.3) |
| /female (ratio) | 87/175 (1:2) | 29/84 (1:3) | 35/50 (2:3) |
| weight (kg) | 116.4 (± 26.1) | 109.9(± 23.6) | 108.3 (± 25.1) |
| kg/m<sup>2</sup>) | 40.4 (± 8.9) | 38.8 (± 7.1) | 38.2 (± 9.1) |
| mg glucose (mg/dL) | 160.78 (± 61.32) | 109.58 (± 15.20) * | 157.08 (± 72.48) |
| c (%) | 7.60 (± 1.50) | 5.91 (± 0.24) * | 7.67 (± 1.77) |
| A-IR | 11.8 (± 13.1) | 7.1 (± 7.4) * | 13.7 (± 17.8) |
| sensitivity C-reactive protein (nmol/L) | 9.31 (± 19.31) | 7.46 (± 7.51) | 9.34 (± 9.10) |
| hydroxybutyrate (mmol/L) | 0.17 (± 0.15) | 0.14 (± 0.13) | 0.15 (± 0.12) |
| il PSQI Score | 7.72 (± 3.72) | 7.96(± 3.43) | 7.92 (± 3.85) |
| ctive sleep quality | 1.18 (± 0.75) | 1.22 (± 0.73) | 1.25 (± 0.79) |
| latency | 1.09 (± 0.93) | 1.33 (± 0.958) | 1.05 (± 0.89) |
| duration | 1.23 (± 0.92) | 1.27 (± 0.96) | 1.14 (± 0.94) |
| ual sleep efficiency | 0.68 (± 0.99) | 0.61 (± 0.89) | 0.71 (± 1.04) |
| disturbances | 1.64 (± 0.63) | 1.66 (± 0.68) | 1.75 (± 0.74) |
| f sleep medication | 0.69 (± 1.16) | 0.66 (± 1.11) | 0.85 (± 1.26) |
| me dysfunction | 1.22 (± 0.77) | 1.21 (± 0.76) | 1.17 (± 0.86) |
| sleepers N (%) | 179 (68.3) | 88 (77.9) | 58 (68.2) |
| sleepers N (%) | 83 (31.7) | 25 (22.1) | 27 (31.8) |

Subjective sleep quality, component 1; sleep latency, component 2; sleep duration, component 3; habitual sleep efficiency, compc disturbances, component 5; use of sleep medication, component 6, and daytime dysfunction, component 7
ue <0.001

### Effect of intervention on sleep Global PSQI and component scores

Overall sleep quality as assessed by the global PSQI score, improved in CCI T2D (median change from 7 to 6; p<0.001) and prediabetes (median change from 7 to 5; p<0.001) groups after one year of the intervention (Figure 1). No significant change in the global PSQI score was observed in UC T2D (median change from 7 to 8, p=0.245). At one year, global PSQI scores in the CCI T2D (p<0.001) and prediabetes (p<0.01) were significantly lower than in the UC T2D, whereas no differences were observed at baseline (Figure 2A). Among patients characterized as poor sleepers at baseline (global PSQI >5), one year global PSQI score was lower in the CCI T2D (p<0.001) and prediabetes (p<0.001) than in the UC T2D (Figure 2B). Greater reduction in the global PSQI score was observed in CCI T2D (median change of −1, p<0.01) and CCI prediabetes groups (median change of −2, p<0.001) compared to the UC T2D group (Figure 3). Further assessment of the PSQI component scores revealed three of the seven components showed significant change at one year for CCI T2D and prediabetes groups. Subjective sleep quality (p<0.01 CCI T2D; p<0.001 CCI prediabetes), sleep disturbance (p<0.01 CCI T2D; p<0.001 CCI prediabetes) and daytime dysfunction (p<0.001 CCI T2D; p<0.001 CCI prediabetes) score were lower in the CCI T2D and prediabetes patients compared to the UC T2D group at one year (Figure 4 A-C).

**Figure 1.**
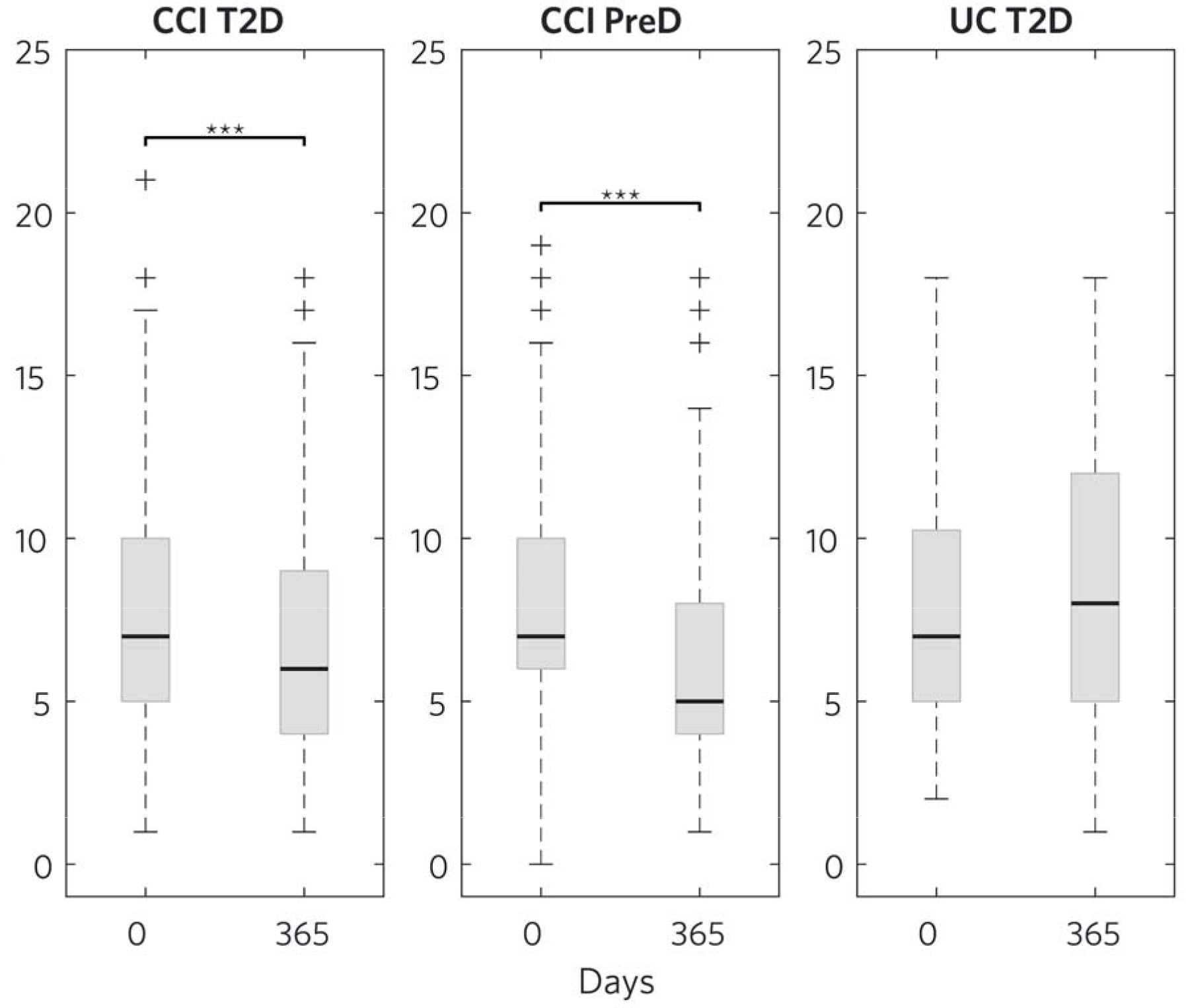
Distribution of global PSQI scores at baseline and 365 days in CCI T2D, CCI PreD and UC T2D. Global PSQI sc icantly reduced in the CCI T2D and CCI PreD groups but not in the UC T2D group after 365 days.

**Figure 2.**
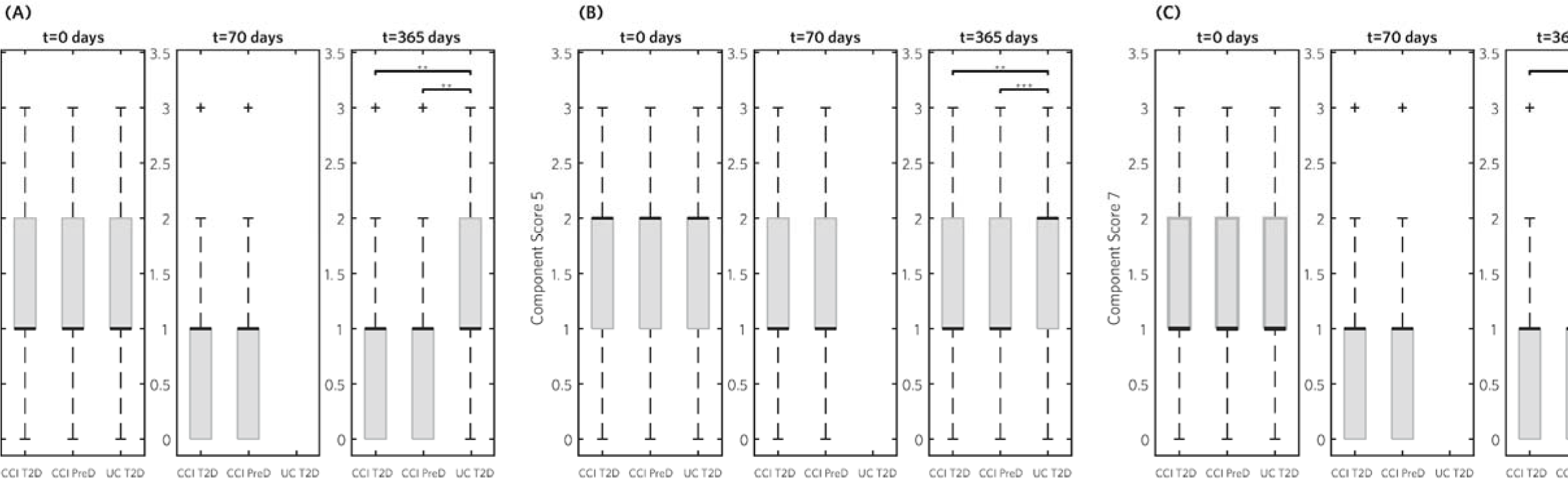
Distribution of PSQI components subjective sleep quality, sleep disturbances and daytime dysfunction in CCI T and UC T2D groups at three different timepoints (0, 70 and 365 days). Subjective sleep quality (A), sleep disturbances ne dysfunction (C) were significantly lower in the CCI T2D and CCI PreD groups when compared to UC T2D group at 36 *‘of descriptors (Figures 1-4) Horizontal line within the box indicates median; upper and lower boundaries of the box n 5th and 75^th^ percentiles; whiskers of the box is the highest and lowest values and “+++” signs represent outlier values. alue <0.01; ***p-value <0.001*

### Resolution of poor sleep quality

There were 179 (68.3%) T2D and 88 (77.9%) prediabetes patients categorized as “poor sleepers” in the CCI at baseline. The proportions of “poor sleepers” in the CCI were reduced after one year of the intervention, with 56.5% of T2D (p=0.001) and 48.7% (p<0.001) of prediabetes patients categorized as “poor sleepers” at one year. In the UC cohort, the proportion of patients categorized as “poor sleepers” did not change after one year (68.2% at baseline to 69.4% at one year).

### Association within the CCI group between changes in global PSQI with metabolic and inflammatory markers

Table 2 shows correlations between changes in the global PSQI score with changes in glucose-related, ketone and inflammatory markers in the CCI. In the prediabetes group, changes in fasting glucose (r= 0.23, p=0.02) and HOMA-IR (r= 0.32, p<0.001) were correlated to changes in PSQI scores after controlling for baseline age, sex and weight. Increased ketone concentrations in the prediabetes participants were also associated with reduction of global PSQI scores (r= −0.242, p=0.01). These correlations observed in the prediabetes group were not present in the CCI T2D group and changes in the HbA1c and hsCRP did not correlate with changes in global PSQI scores in either group. Change in mean weight (p=0.04) and HOMA-IR (p=0.01) were the only variables independently and significantly associated between the three different sleep status (improved, maintained and not improved sleep status) at one year of the intervention. No statistically significant differences were found in weight loss changes between patients with improved, maintained and not improved sleep status. Patients who maintained sleep showed highest reductions of HOMA-IR (−6.94 ± 0.86), with statistically significant difference than those who did not improve sleep, after one year of the intervention (p = 0.02). Improvements in HOMA-IR among patients in the improved sleep (−4.17 ± 0.86) and not improved sleep status (−4.24 ± 0.55) did not differ significantly.

**Table 2.** Correlation analyses between change in the global PSQI score and change in metabolic parameters after one year of rmari and ^+^adjusted Pearson’s correlations. Adjustments while controlling for age, sex and baseline weight

|  | CCI T2D Cohort |  |  |  | CCI Prediabetes Cohort |  |  |  |
| --- | --- | --- | --- | --- | --- | --- | --- | --- |
|  | N=262 |  |  |  | N=113 |  |  |  |
|  | *rho* | *P*<br>*value* * | Adjusted *r* | *P*<br>*value*<sup>+</sup> | *rho* | *P*<br>*value* * | Adjusted *r* | *P value*<sup>+</sup> |
| glucose (mg/dl) | 0.032 | 0.60 | 0.008 | 0.90 | 0.240 | 0.01 | 0.226 | 0.018 |
| %) | -0.037 | 0.55 | -0.049 | 0.44 | -0.024 | 0.80 | -0.032 | 0.74 |
| R | -0.060 | 0.34 | -0.069 | 0.27 | 0.314 | 0.0008 | 0.323 | 0.0006 |
|  | -0.003 | 0.96 | -0.044 | 0.49 | -0.297 | 0.002 | -0.242 | 0.011 |
|  | -0.067 | 0.29 | -0.008 | 0.90 | -0.022 | 0.82 | -0.032 | 0.74 |

### Effect of persistent pain on sleep improvement

We further assessed the effect of pain on sleep improvement in the CCI by classifying the patient’s pain status using response retrieved from questions specifically related to pain in the sleep and knee (KOOS) questionnaires. As illustrated in supplementary figure 2, patients with pain had higher global PSQI scores, indicating poorer sleep, compared to those categorized under “non-pain” group at all three time points. Both patients in the “non-pain” (Supplementary figure 3A, p<0.001). and “pain” group (Supplementary figure 3B, p<0.01) had reductions in their global PSQI score at 70 days and one year.

### Effect of shifted sleep chronotype on sleep improvement

We also assessed the effect of shifted sleep chronotype on the global PSQI score improvement. Patients were classified as having shifted sleep chronotype based on their self-reported wake-up times and bedtimes as defined in the methods. There were 18, 27, and 96 patients in the CCI cohort classified as both wake-up time and bedtime shifted, wake-up time shifted only or bedtime shifted only respectively. Patients with shifted bedtimes, had reduced global PSQI scores (p<0.01), as did those with normal chronotype (p<0.001) (Supplementary figures 4A and B). However, those patients with shifted wake-up times (Supplementary figures 4C) did not show a change in their global PSQI score after one year of the intervention. Those with both shifted wake-up times and bedtimes also did not show a change in their global PSQI score after one year of the intervention.

### Effect of CPAP usage on sleep improvement

At baseline, there were a total of 140 participants in both CCI and UC treatment groups with CPAP equipment prescribed for sleep. Among CPAP users, 91 were in the CCI T2D group, 31 in the CCI prediabetes and 18 in the UC T2D group. Fifteen (13 CCI T2D and 2 UC T2D) of the 140 participants discontinued using CPAP at one year. Only 6 (46%) of the 13 CCI T2D participants discontinued due to patient-reported improvement in sleep quality from the CCI and reduction of weight; the remaining 7 reported dis-continuation due to discomfort or personal choice. Global PSQI scores among the CPAP users at baseline and one year did not show a significantly different distribution pattern than what was observed in the full cohort of participants.

## Discussion

This study is one of the first designed to assess the effect of carbohydrate restriction and nutritional ketosis on sleep quality in individuals with hyperglycemia and insulin resistance. Improved patient-reported sleep quality as assessed by global PSQI suggests that CCI including nutritional ketosis benefited sleep quality in both patients with T2D and prediabetes. The proportion of patients categorized as “poor sleepers” at one year was significantly reduced in the CCI groups but not in the UC group. Furthermore, these results demonstrate that the sleep quality improvement observed in the whole intervention population was due in part to 17% of baseline “poor sleepers” being reclassified as “good sleepers” at one year. Our results are consistent with previous findings that showed improved overall sleep quality in children consuming ketogenic diets ^31, 32^.

Improvement in the global PSQI score of patients undergoing the CCI was mainly due to significant changes in three PSQI components: subjective sleep quality, sleep disturbance and daytime dysfunction. Both objective and subjective sleep quality impairment are frequently reported in diabetes patients and positively associated with severity of hyperglycemia ^8-11^. Likewise, correlation between poor sleep quality and increased carbohydrate intake ^30^ is also previously reported. These observed patterns of association between sleep quality with hyperglycemia and carbohydrate intake may explain why this carbohydrate restriction intervention improved subjective sleep quality. The sleep disturbance component of the global PSQI score is associated with poor glycemic control among T2D patients ^43^. One study reported a significant correlation between sleep disturbance and HbA1c level ^44^. Night time sleep disturbance in T2D patients can be related to a wide range of conditions such as nocturnal polyuria, pain, and breathing problems, especially in those with OSA. In our study, we also showed that patients encountering persistent pain, including knee pain, had a higher median global PSQI score, while one year of the intervention effectively improved global PSQI scores in these patients despite the persistence of reported pain in some patients. It is possible that improvement in the sleep disturbance of the CCI patients contributed to the glycemic control improvement in these patients. The effectiveness of the intervention in improving sleep in those with pain, further emphasizes its’ applicability in alleviating sleep disturbance.

Furthermore, there was a significant improvement in the daytime dysfunction component of the global PSQI score in the CCI group. Excessive daytime sleepiness and dysfunction are reported commonly in T2D ^45, 46^, and weight loss through bariatric surgery has a positive resolving effect on daytime dysfunction and sleepiness ^47, 48^. In the present investigation, the majority of CCI patients achieved weight loss of ≥ 10%, which could have contributed to the significant improvement observed in daytime function. In addition, we also evaluated the effect of the intervention on a subcohort of patients with a self-reported pattern of shifted non-standard bedtimes and wake-up times that were not aligned to the light dark cycle, which likely affects daytime functioning. Circadian rhythm disruption is frequently associated with metabolic alterations, especially in an insulin resistant state ^49, 50^. While patients with a normal sleep chronotype benefited the most, the intervention also improved the sleep of patients with time shifted bedtimes. A similar advantage of the intervention was not observed in patients with shifted wake-up times, though this may be due to the limited number of patients in this subgroup (n=27).

The improvement in the global PSQI score observed in CCI patients occurred concurrently with weight reduction and glycemic control improvement ^34, 35^. Martin et al ^51^ reported a direct correlation between degree of weight loss and global PSQI score improvement in healthy nonobese adults receiving an energy restricted diet, while Chaput et al^52^ reported an improvement in global PSQI score following the initial 5-kg weight loss, but no additional improvement with subsequent weight loss. A study using a ketogenic diet in children alleviated abnormal sleep architecture; however, weight loss was suggested as the main determinant of improved sleep ^33^. These studies collectively imply a direct association between weight loss and improved PSQI score. Likewise, long-term maintenance of weight loss was associated with better sleep quality and quantity (); while the degree of weight loss reduction is directly correlated with OSA improvement(). However, some studies also demonstrate the efficacy of anti-glycemic medications for improving PSQI score concurrent with improved glycemic control ^53^. This study identified associations between HOMA-IR and weight reductions with stratification of patients’ sleep status in the full CCI cohort even though there were no significant differences in weight loss and insulin resistance reduction levels between those who had improved sleep and those who did not. Patients with good sleep quality at the beginning of the intervention benefited the most in reducing insulin resistance. Improvement in fasting glucose and HOMA-IR were only positively associated with improved PSQI score in prediabetes patients.

It is not clear if nutritional ketosis achieved by substantial carbohydrate restriction augmented the effect of the intervention on sleep or if weight loss and/or improved glycemic control generated from the intervention contributed to sleep quality improvements. We showed a significant correlation between blood beta-hydroxybutryrate (BHB) levels and PSQI improvement in the prediabetes cohort. While the effect of and mechanism of BHB in sleep are not clear, a positive correlation between blood BHB levels and carbon dioxide (CO_2_) response was previously reported in patients with obesity related hypoventilation syndrome that had reduced CO_2_ response ^54^. A continuous state of ketosis through carbohydrate restriction and fat intake also induces the postprandial release of a satiety hormone, cholecystokinin (CCK)^28, 55, 56^. When administered in rats, CCK was shown to promote slow wave activity and NREM sleep ^57^. CCK was also shown to induce sleep when administered in diabetic rats ^58^. Therefore, it is possible that one mechanism of improved sleep with a ketogenic diet that increases BHB levels is through CCK induction.

There are several limitations of our study. The study was designed mainly to assess the impact of the CCI on glycemic control, medication use, weight, and cardiovascular disease risk factors. Patient-reported outcomes for quality of life measures including sleep were included as secondary endpoints. It is difficult to determine the causality among the intervention, improvement in primary outcomes and improvement in sleep from this study. A major limitation of this study is the use of subjective sleep measures as self-reported sleep assessment is subject to limited self-knowledge of sleep behavior and inconsistency in reporting. Therefore, future studies that use randomized controlled trial designs and objective sleep measures are needed to confirm our results. In addition, patients with an established diagnosis of a sleep disorder such as OSA were not separated in the analysis since complete records of their CPAP usage were not collected in the questionnaire. Patient compliance with CPAP usage is essential for making interpretations about the status of their OSA treatment and its effect on sleep and glycemic control. The study also lacked recruitment of prediabetes patients in the UC group for direct comparison of the treatment effect between UC and CCI on sleep in these patients.

In conclusion, these results demonstrate that overall sleep quality significantly improved in T2D and prediabetes patients undergoing remote CCI including nutritional ketosis but not in T2D patients in the UC group. The sleep improvement was concurrent with weight reduction and glycemic control improvement. The PSQI components that improved were sleep quality, sleep disturbance and daytime dysfunction. These results suggest that nutritional ketosis benefits overall health through improved glycemic control as well as improved sleep quality.

## Supporting information

**Author contributions**

S.J.A, M.S, C.J.V and J.P.M drafted the manuscript. A.L.M, N.H.B, S.J.H and S.J.A participated in data acquisition and compiling. M.S and S.J.A analyzed the data. C.J.V supervised this particular analysis, J.P.M, A.L.M, S.J.H, N.H.B, W.W.C, S.D.P and J.S.D edited the manuscript. W.W.C. proposed measuring subjective sleep quality as part of the parent Continuous Care Intervention clinical trial. All authors approved the final version of the manuscript.

## Abbreviations

CCI: continuous care intervention
UC: usual care
T2D: type 2 diabetes
BMI: body mass idex
PSQI: Pittsburgh Sleep Quality Index
OSA: obstructive sleep apnea
HbA1c: hemoglobin A1c
CPAP: continuous positive airway pressure
AHI: apnea and hypopnea indices
KD: ketogenic diet
REM: rapid eye movement
SWS: slow wave sleep
BHB: beta-hydroxybutryrate
HOMA- IR: homeostatic model assessment of insulin resistance
hsCRP: high-sensitivity C-reactive protein

