## Supplementary material for "Improvement in Patient-Reported Sleep in Type 2 Diabetes and Prediabetes Participants Receiving a Continuous Care Intervention with Nutritional Ketosis"

**Supplementary Figure Legends**

**Supplementary Figure 1**. Individual changes in PSQI over 1 year (A) CCI T2D (B) CCI Prediabetes (C) UC T2D

**Supplementary Figure 2**. Distribution of global PSQI scores in CCI T2D, CCI PreD and UC T2D at three different timepoints (0, 70 and 365 days) (A) Among the full patient cohort, global PSQI were significantly lower in the CCI T2D and CCI PreD when compared to UC T2D at 365 days. (B) Among the “poor sleepers” at baseline, global PSQI were significantly lower in the CCI T2D and CCI PreD when compared to UC T2D after 365 days. (UC T2D patients were not surveyed at 70 days.)

**Supplementary Figure 3.** Distribution of change in global PSQI score in CCI T2D, CCI PreD and UC T2D after 365 days. The scores showed significant reduction in the CCI T2D and CCI PreD groups relative to baseline and to the UC T2D group.

**Supplementary Figure 4.** Distribution of global PSQI scores in “non-pain” and “pain” patients among the CCI at three different timepoints (0, 70 and 365 days)

**Supplementary Figure 5.** Distribution of global PSQI score in (A) “non-pain” and (B) “pain” patients among the CCI after 365 days. The scores showed significant reduction in the CCI “non-pain” and “pain” groups relative to baseline

**Supplementary Figure 6.** Distribution of global PSQI in (A) “Shifted bedtime” (B) “Normal bedtime” (C) “Shifted wake-up time” (D) “Normal wake-up time” patients among the CCI after 365 days. The scores showed significant reduction in the CCI “Shifted bedtime”, “Normal bedtime” and “Normal wake time” groups relative to baseline

*Boxplot descriptors (Figures 2-4) Horizontal line within the box indicates median; upper and lower boundaries of the box represent the 25th and 75th quartiles; whiskers of the box is the highest and lowest values and “+++” signs represent outlier values.*

*** p-value <0.01; *** p-value <0.001*

**Supplementary Figure 1**

A	B	C
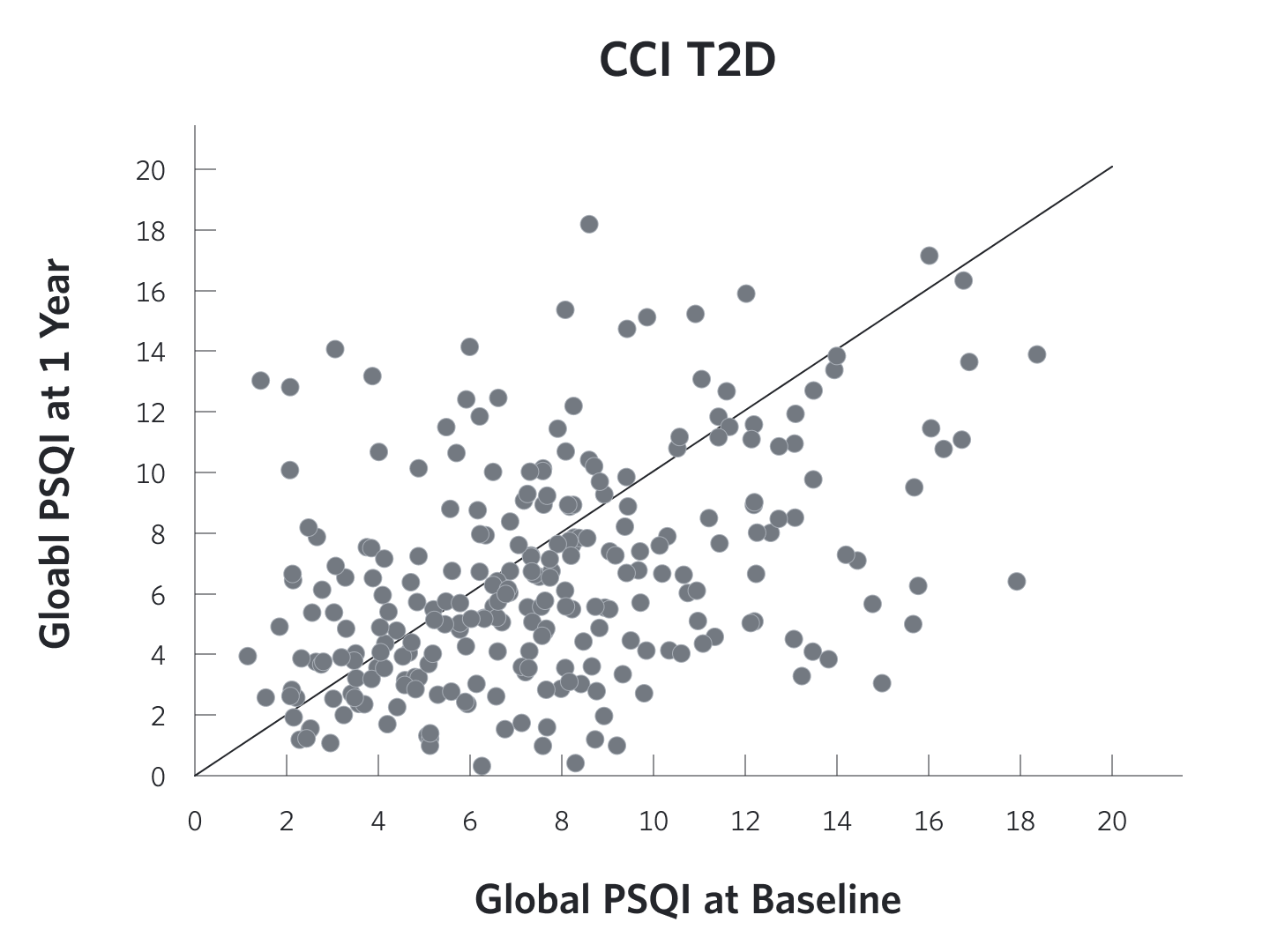

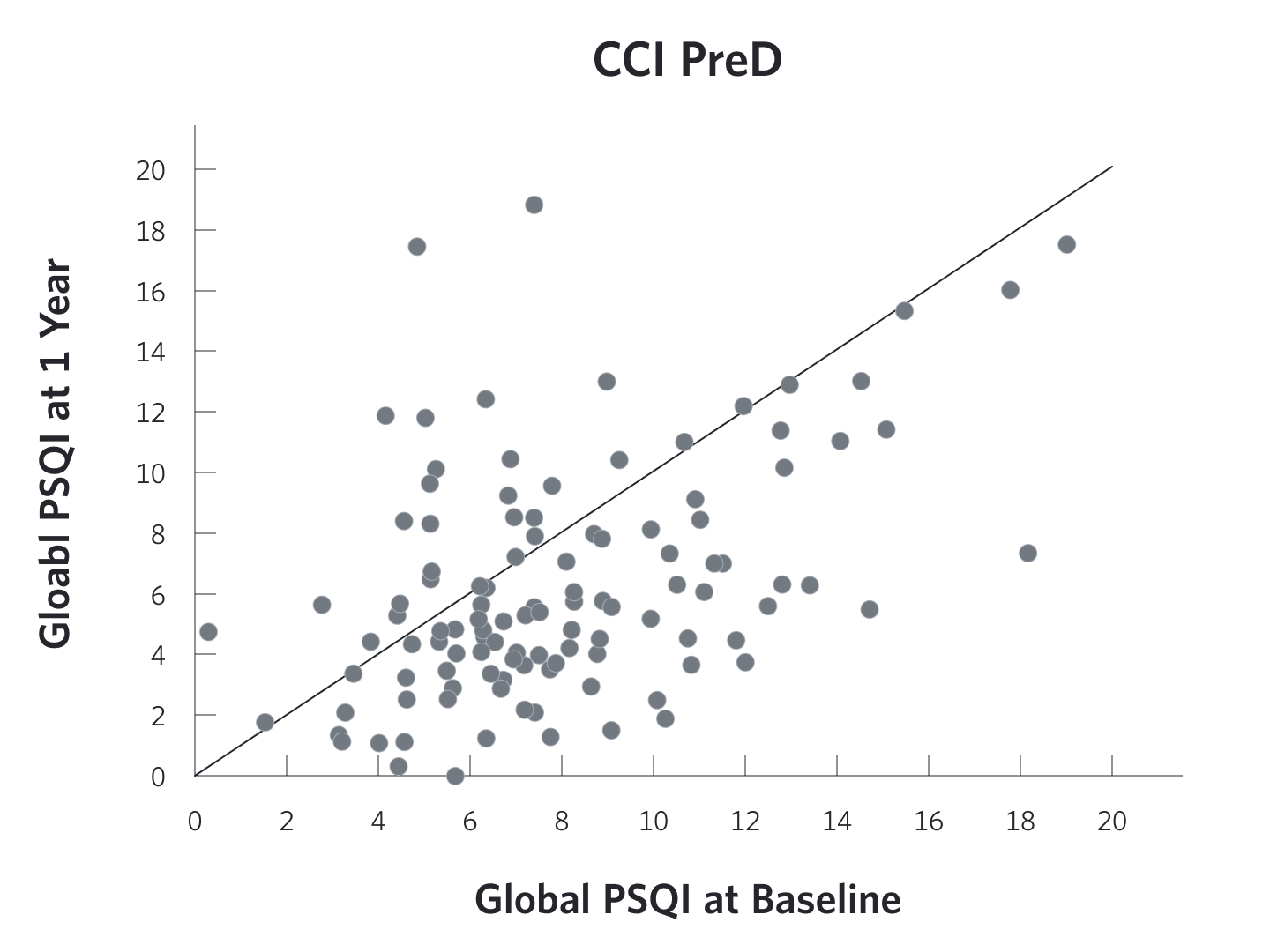

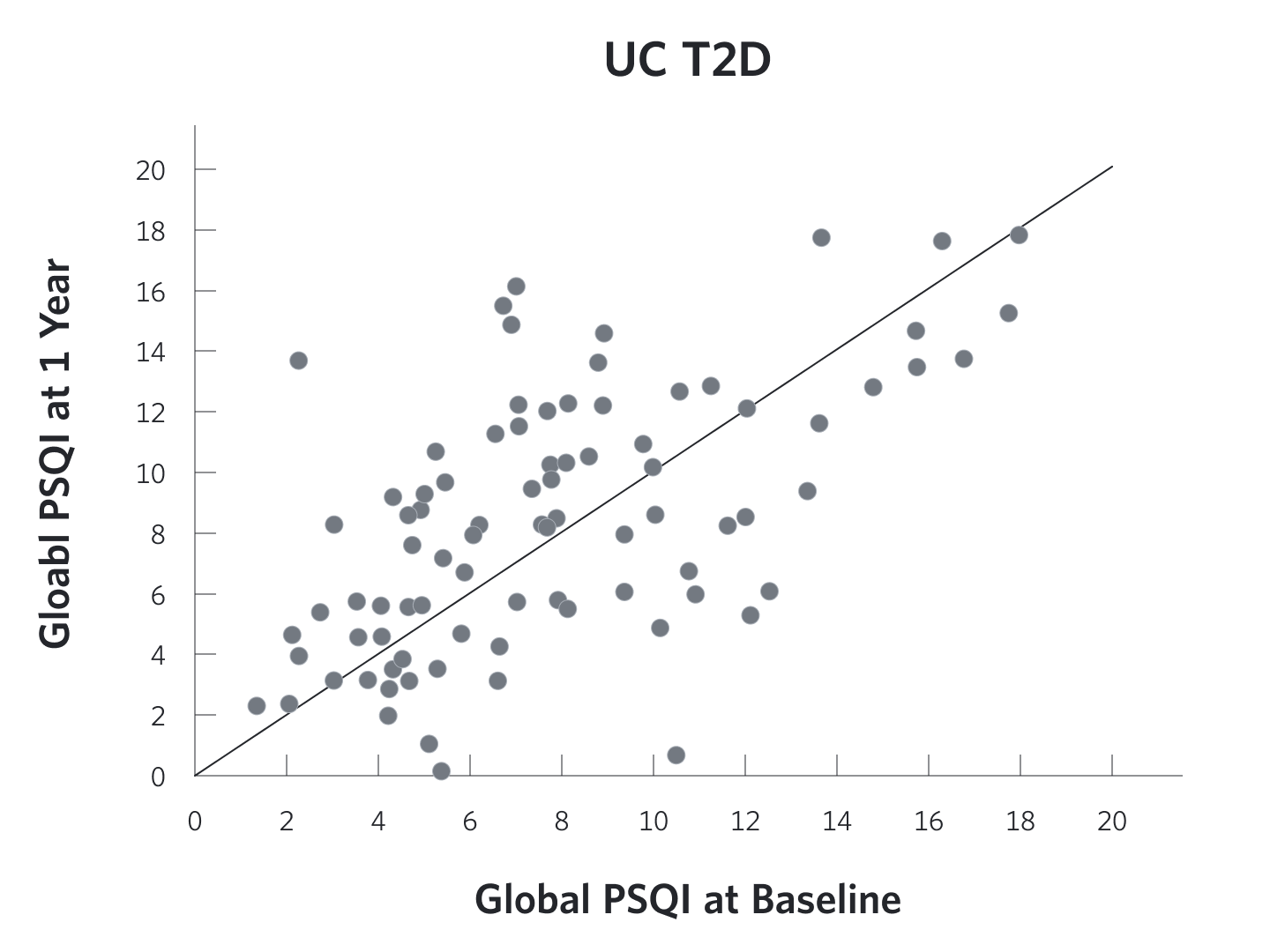

**Supplementary Figure 2**

A	B

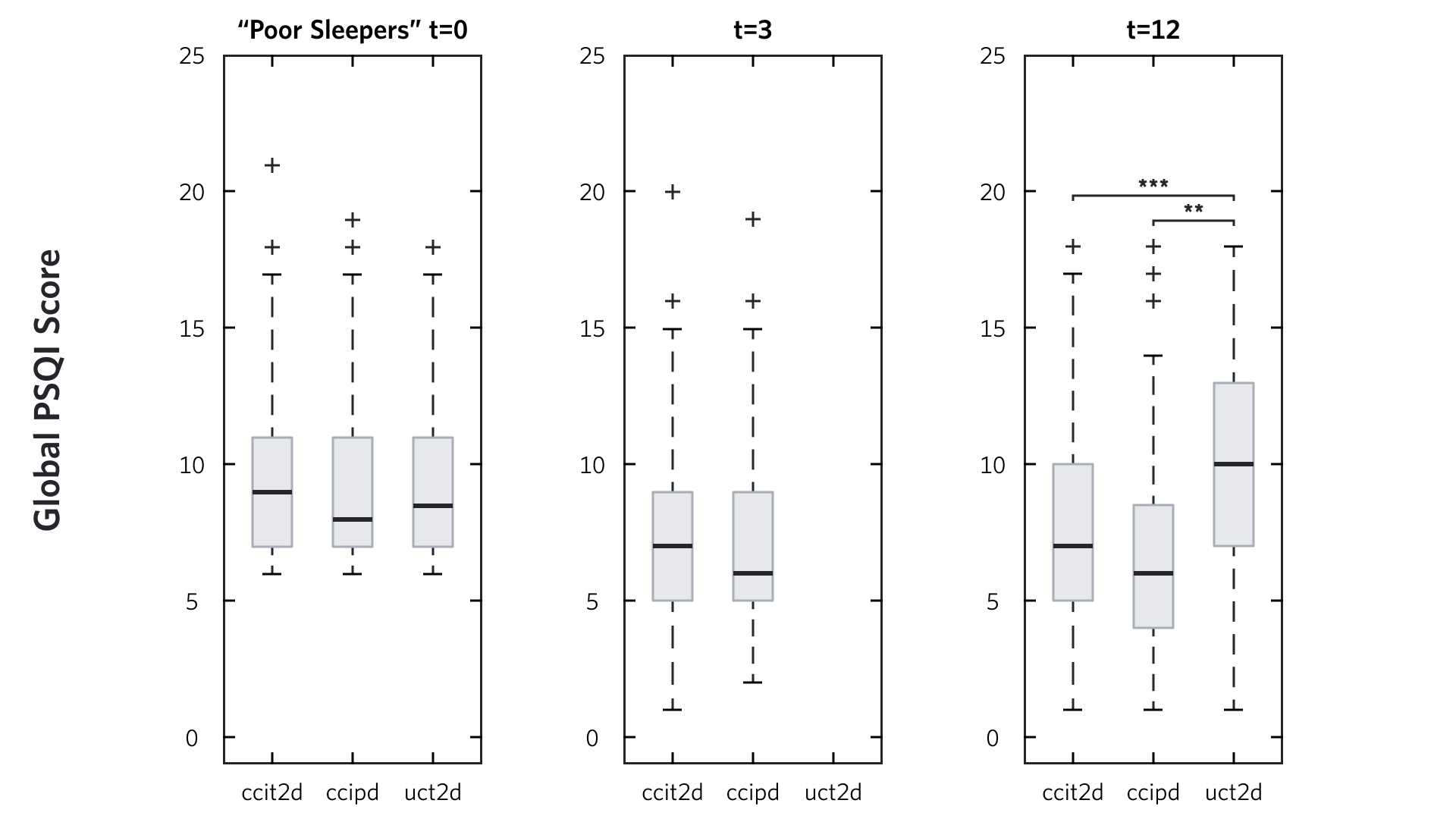

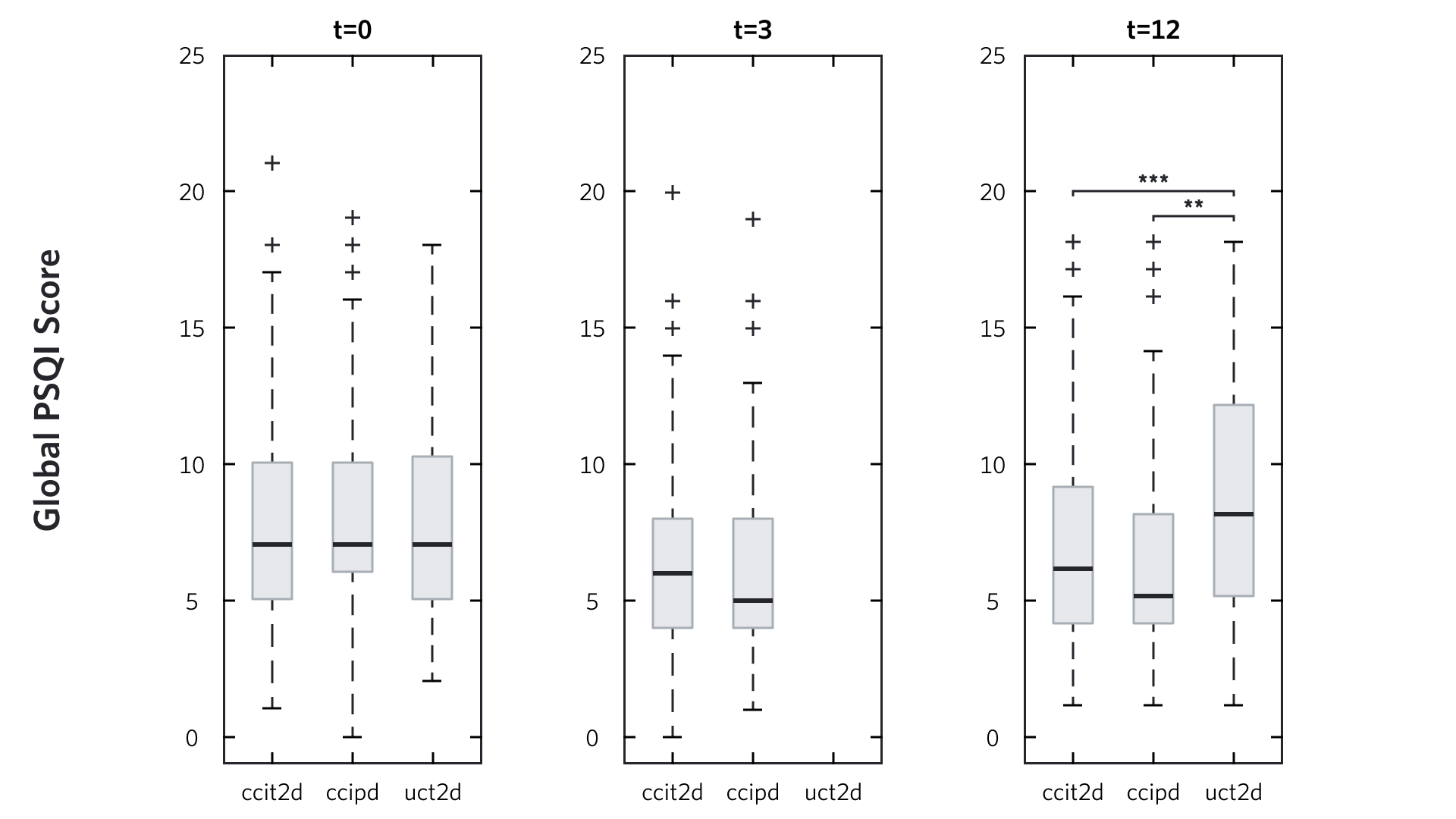

**Supplementary Figure 3**

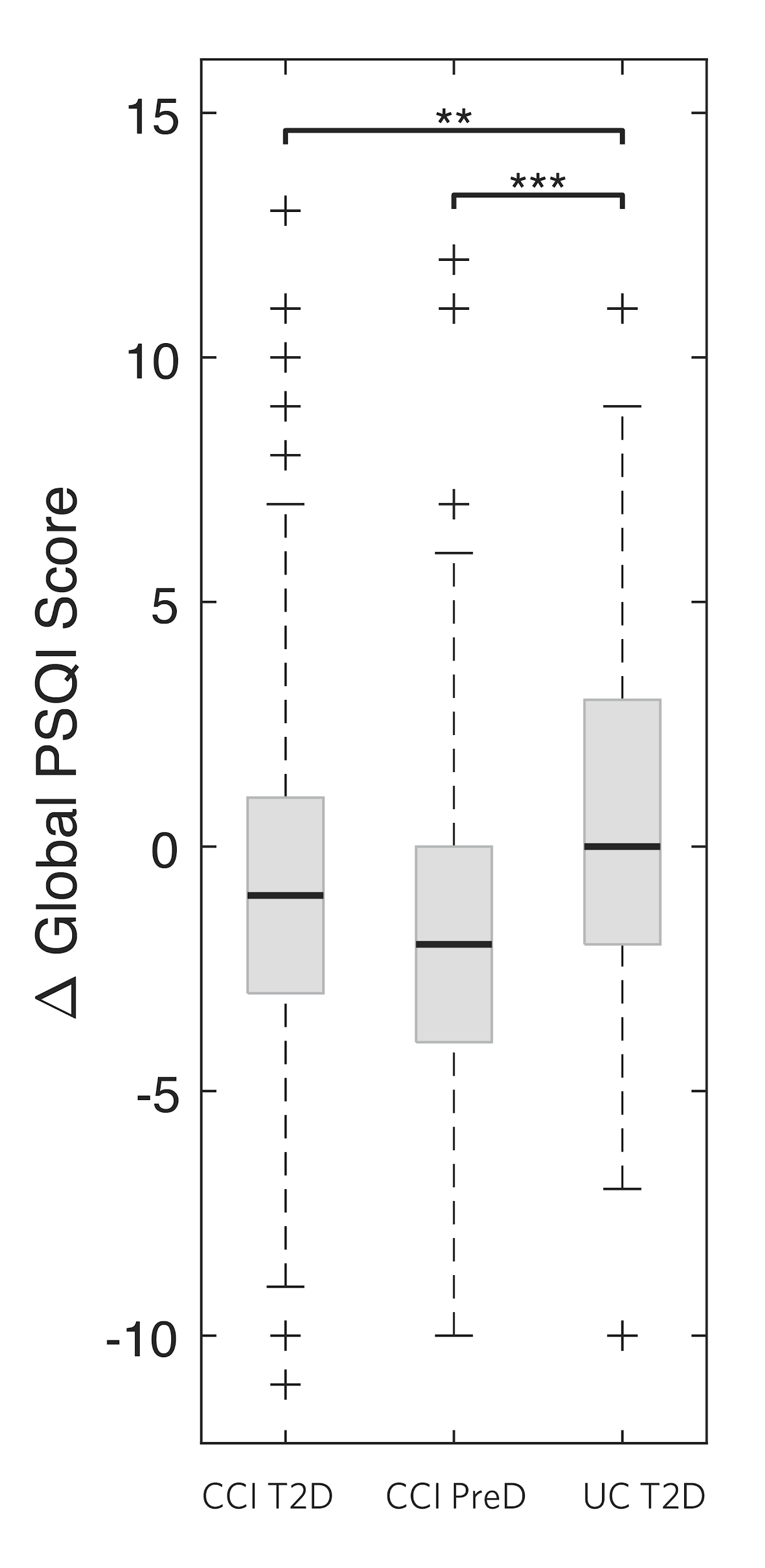

**Supplementary Figure 4**

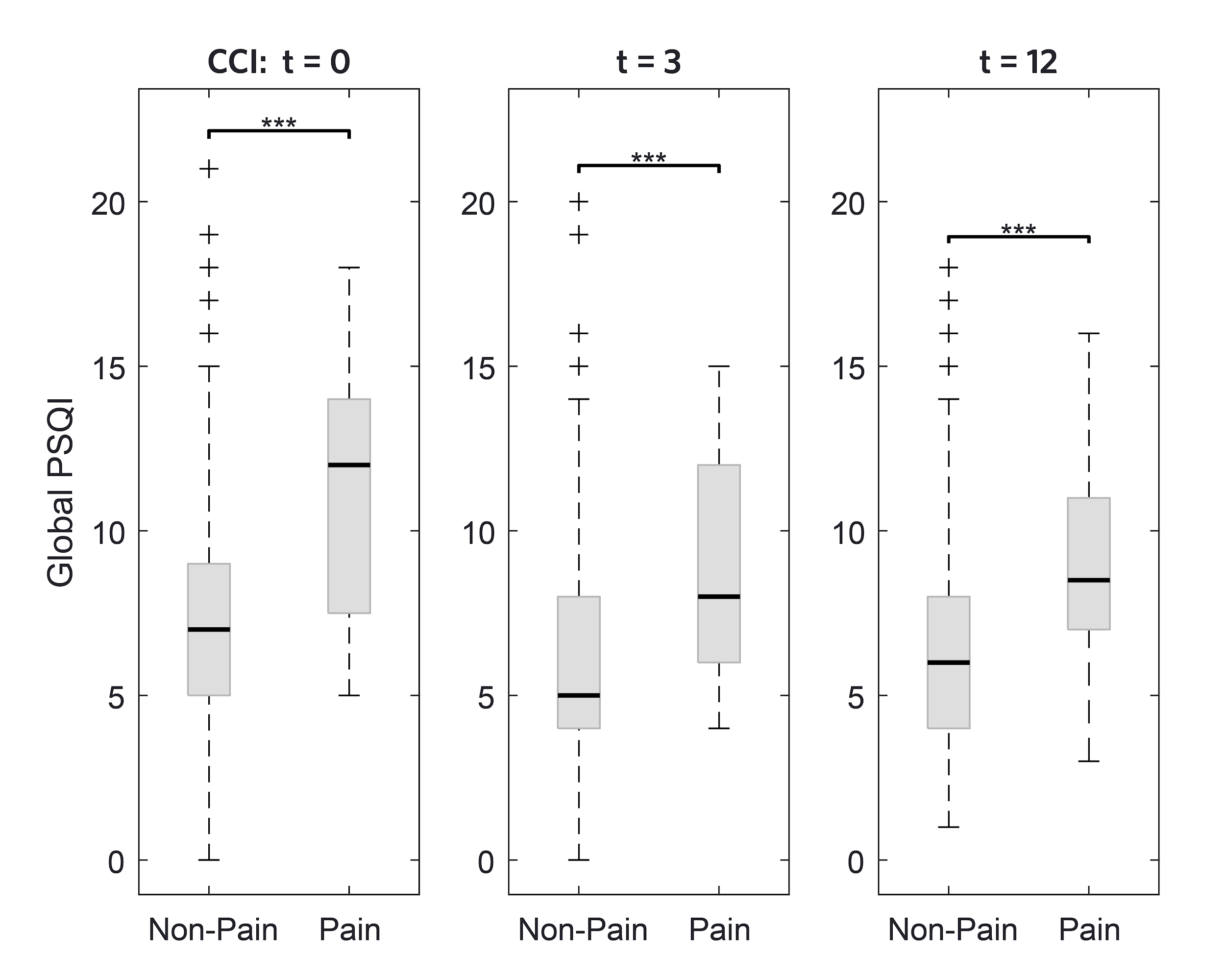

**Supplementary Figure 5**

A	B

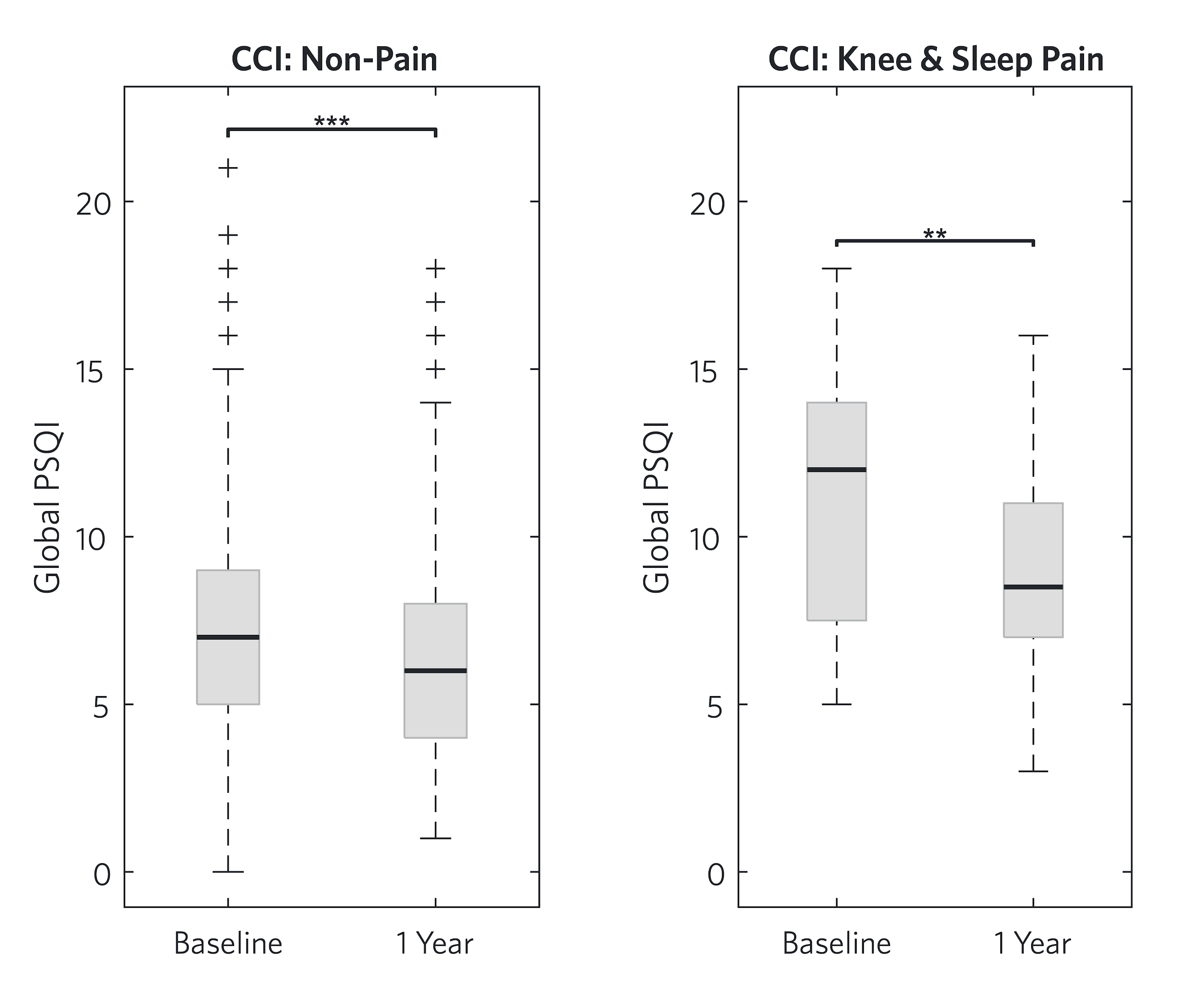

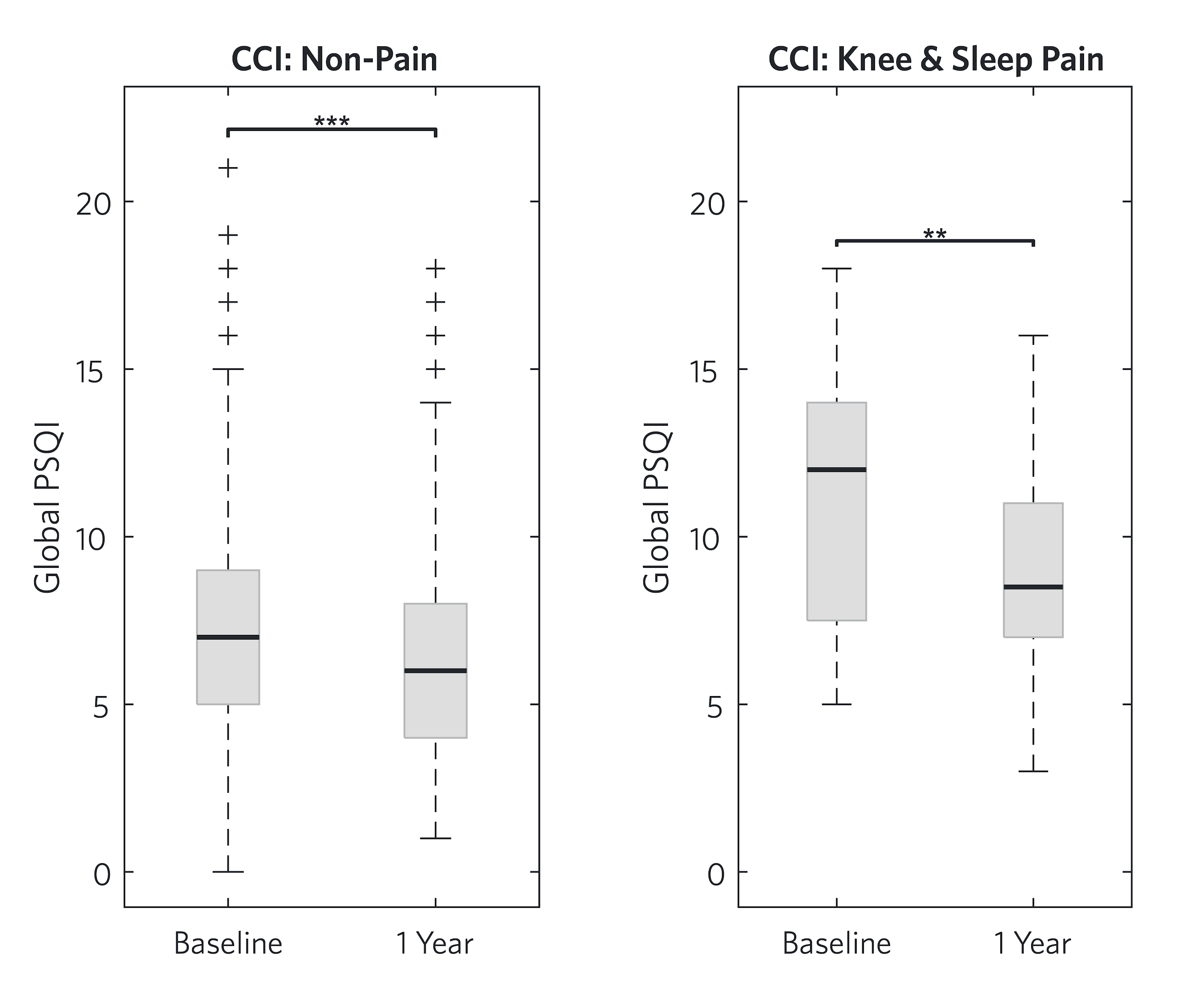

**Supplementary Figure 6**

A	B	C	D

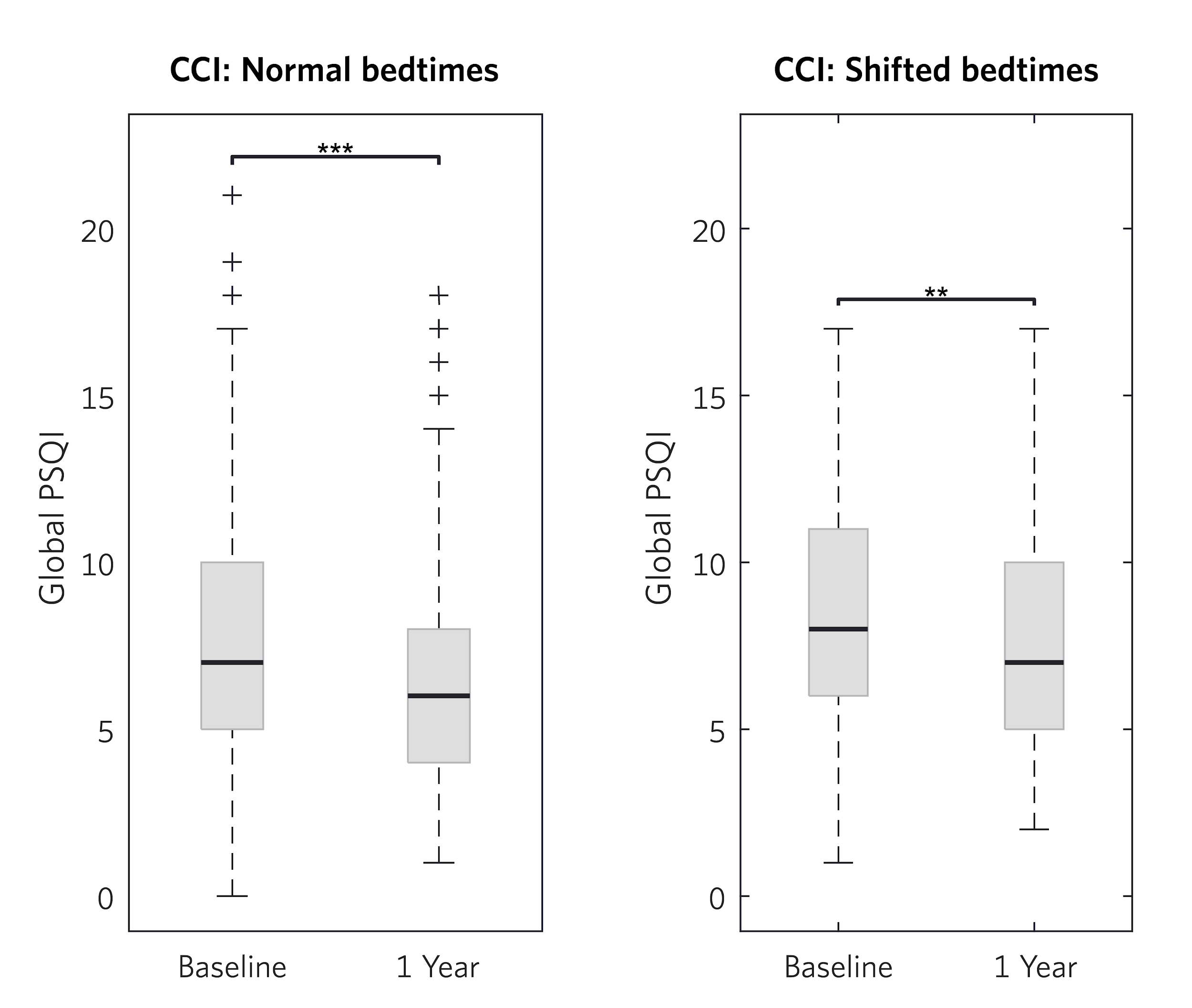

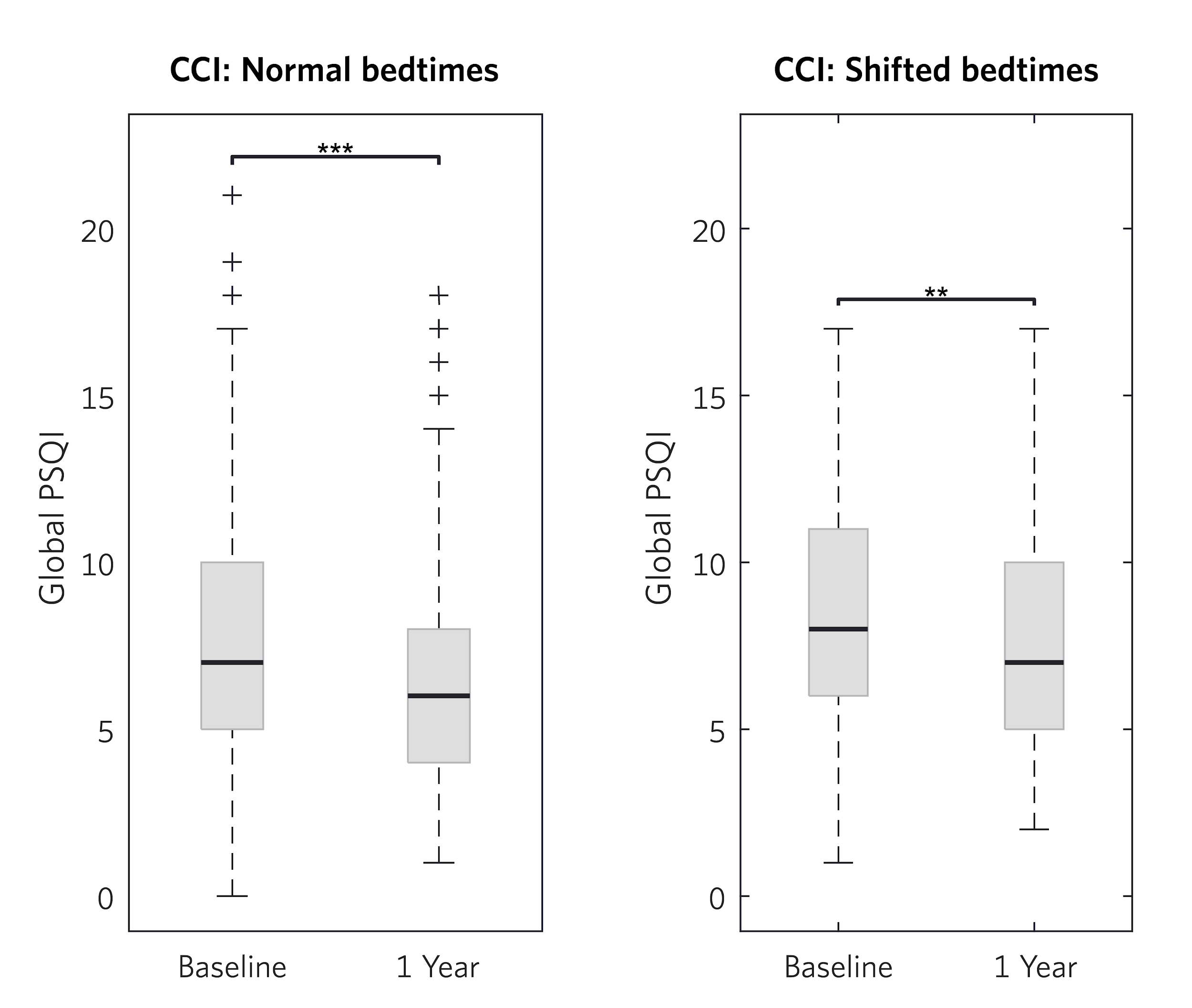

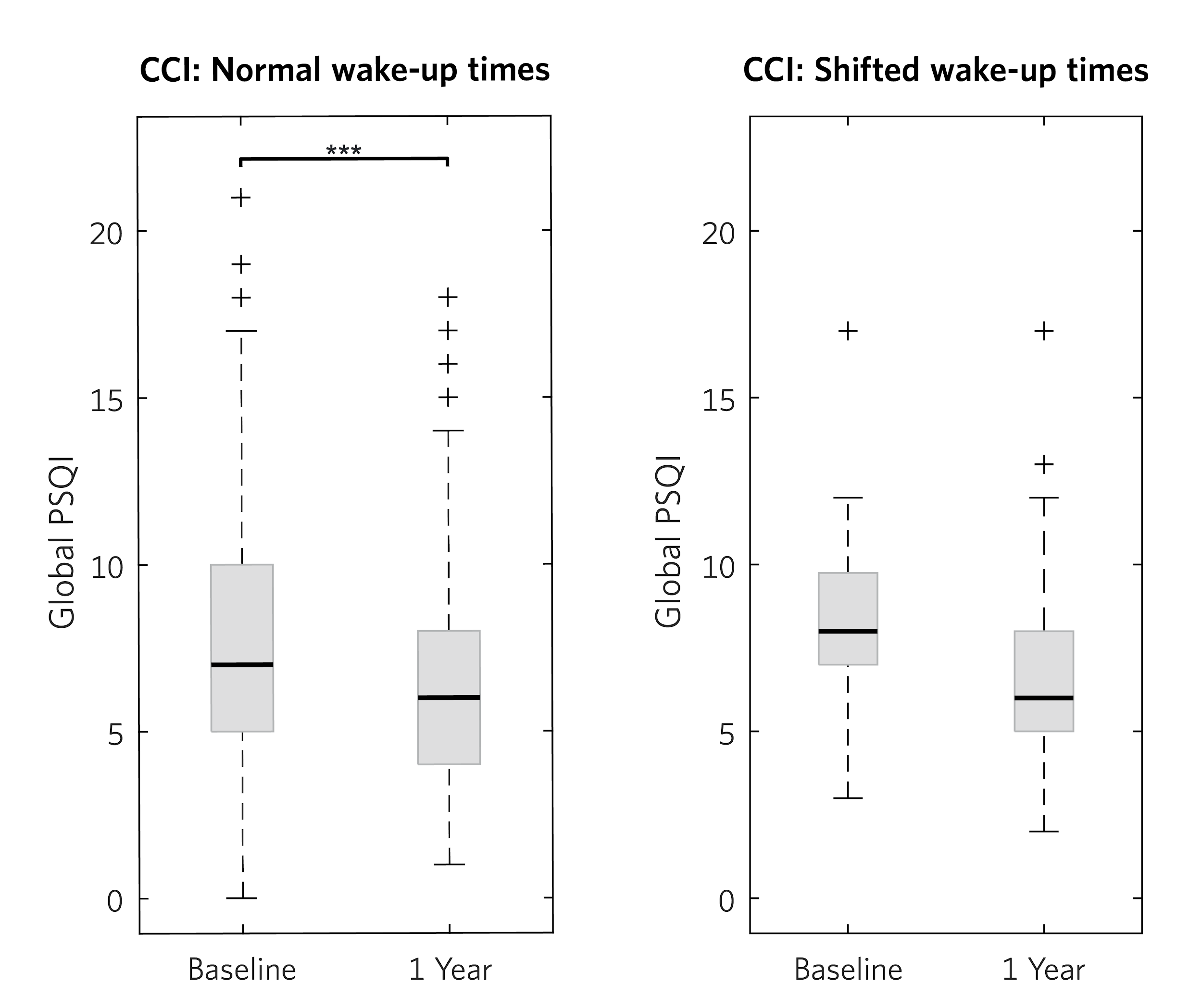

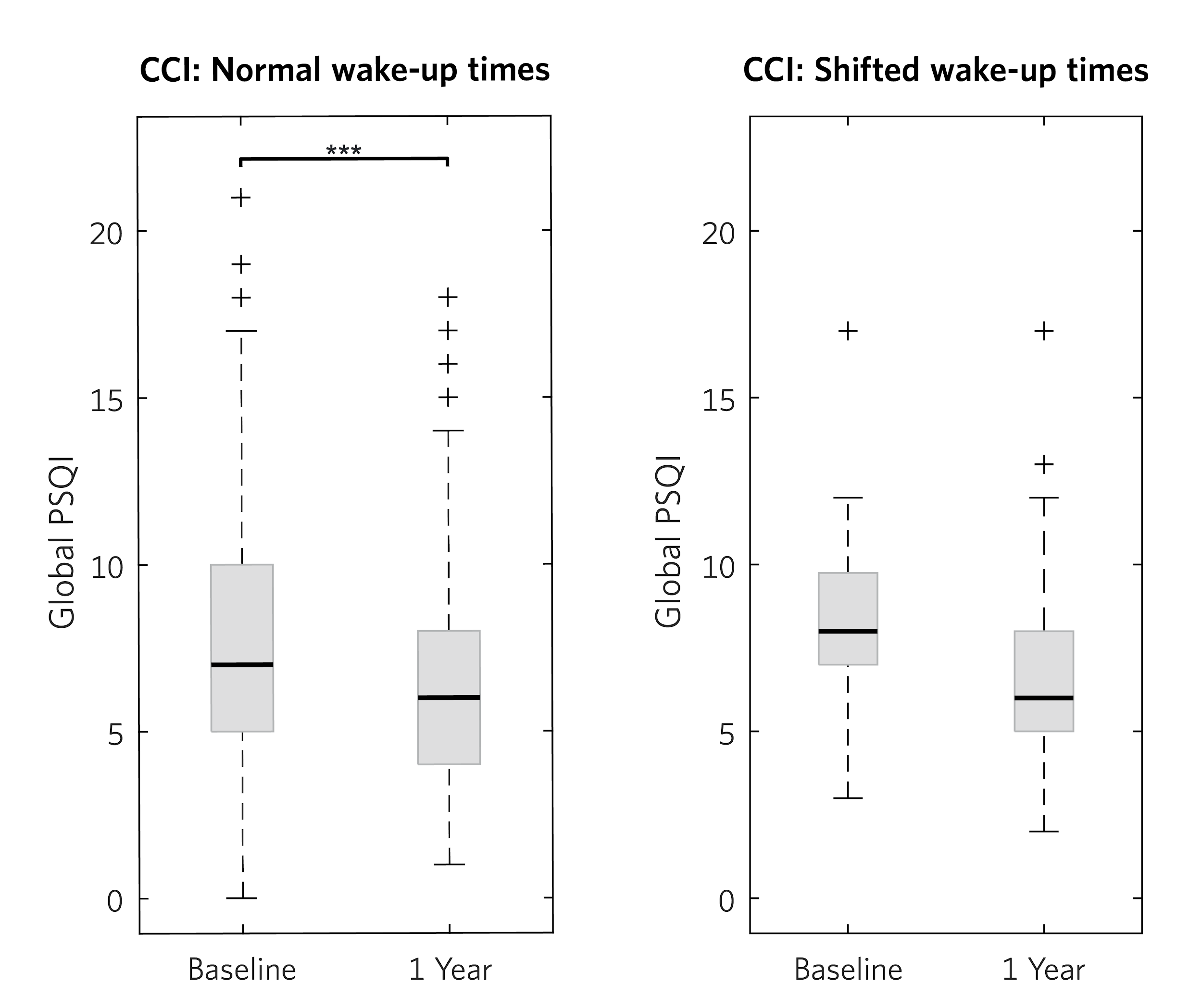
