## Supplementary material for "Improvement in Patient-Reported Sleep in Type 2 Diabetes and Prediabetes Participants Receiving a Continuous Care Intervention with Nutritional Ketosis"

**Supplementary Table 1.** Baseline characteristics of completers and non-completers (CCI Type 2 Diabetes, CCI Prediabetes and UC Type 2 Diabetes)

|  | **All** | | **Completers with PSQI data** | | **Non-completers without PSQI data** | | **Completers-Non Completers** |
| --- | --- | --- | --- | --- | --- | --- | --- |
|  | **N** | **Median (IQR)** | **N** | **Median (IQR)** | **N** | **Median (IQR)** | **P-value** |
| **Age (years)**  CCI Type 2 Diabetes  CCI Prediabetes  UC Type 2 Diabetes | 230 96 66 | 56 (12)  54 (13)  55 (13) | 158 66 52 | 57 (11)  55 (12)  56 (13) | 72 30 14 | 53 (14)  53 (20)  51 (12) | 0.01  0.26  0.53 |
| **Female (%)**  CCI Type 2 Diabetes  CCI Prediabetes  UC Type 2 Diabetes | 230  96  66 | 33.5  25.0  40.9 | 158  66  52 | 36.1  28.8  40.4 | 72  30  14 | 27.8  16.7  42.9 | 0.28  0.31  1.00 |
| **Body weight (kg)**  CCI Type 2 Diabetes  CCI Prediabetes  UC Type 2 Diabetes | 226  95  62 | 113.4 (30.8) 106.0 (29.5) 103.5 (26.9) | 155  65  48 | 110.6 (29.4) 107.9 (34.3) 105.1 (28.9) | 71  30  14 | 116.6 (34.7) 102.5 (22.0)  98.9 (21.7) | 0.68  0.36  0.51 |
| **BMI (kg/m^2^)**  CCI Type 2 Diabetes  CCI Prediabetes  UC Type 2 Diabetes | 230  96  59 | 39.2 (10.6)  37.9 (8.9)  34.7 (10.2) | 158  66  47 | 39.0 (10.6) 37.8 (9.8) 35.3 (10.2) | 72  30  12 | 39.8 (11.5) 37.9 (5.3) 33.7(8.5) | 0.41  0.67  0.14 |
| **Fasting glucose (mg/dL)**  CCI Type 2 Diabetes  CCI Prediabetes  UC Type 2 Diabetes | 227  94  66 | 147.00 (72.00)  107.00 (16.00)  136.00 (68.00) | 156  65  52 | 147.50 (64.00)  107.00 (17.25)  138.50 (69.50) | 71  29  14 | 145.00 (83.50)  107.00 (10.50)  123.00 (61.00) | 0.91  0.91  0.54 |
| **Hemoglobin A1c (%)**  CCI Type 2 Diabetes  CCI Prediabetes  UC Type 2 Diabetes | 230  94  66 | 7.1 (1.7)  5.9 (0.3)  7.2 (2.0) | 158  64  52 | 7.1 (1.6)  5.8 (0.3)  7.2 (2.2) | 72  30  14 | 7.4 (1.9)  5.9 (0.3)  7.0 (1.2) | 0.48  0.22  0.53 |
| **HOMA-IR**  CCI Type 2 Diabetes  CCI Prediabetes  UC Type 2 Diabetes | 213  87  61 | 7.9 (8.9)  5.3 (5.2)  6.3 (9.4) | 148  61  48 | 7.8 (8.0)  5.1 (5.3)  5.9(9.9) | 65  26  13 | 8.4 (12.0)  6.3 (5.6)  10.7 (7.6) | 0.58  0.36  0.36 |
| **hsC-reactive protein (nmol L^-1^)**  CCI Type 2 Diabetes  CCI Prediabetes  UC Type 2 Diabetes | 219  84  65 | 5.2 (8.2)  4.9 (5.9)  5.3 (7.6) | 150  60  52 | 5.3 (8.1)  4.2 (4.2)  4.7 (8.0) | 69  24  13 | 5.2 (8.5)  7.5 (6.9)  6.7 (5.7) | 0.47  0.14  0.23 |
| **Beta-hydroxybutyrate (mmol L^-1^)**  CCI Type 2 Diabetes  CCI Prediabetes  UC Type 2 Diabetes | 216  88  61 | 0.12 (0.12)  0.09 (0.07)  0.10 (0.09) | 150  62  48 | 0.12 (0.12)  0.09 (0.08)  0.09 (0.09) | 66  26  13 | 0.12 (0.12)  0.09 (0.07)  0.13 (0.08) | 0.83  0.87  0.04 |
| **Global PSQI Score**  CCI Type 2 Diabetes  CCI Prediabetes  UC Type 2 Diabetes | 230  96  66 | 7.00 (5.00)  7.00 (4.00)  7.50 (5.00) | 158  66  52 | 7.00 (4.00)  7.00 (4.00)  7.50 (5.00) | 72  30  14 | 8.00 (6.50)  8.00 (5.00)  7.50 (5.00) | 0.47  0.52  0.47 |
| **Subjective sleep quality**  CCI Type 2 Diabetes  CCI Prediabetes  UC Type 2 Diabetes | 230  96  66 | 1.00 (1.00)  1.00 (1.00)  1.00 (1.00) | 158  66  52 | 1.00 (0.00)  1.00 (1.00)  1.00 (0.50) | 72  30  14 | 1.00 (1.00)  1.00 (1.00)  1.00 (1.00) | 0.43  0.70  0.80 |
| **Sleep latency**  CCI Type 2 Diabetes  CCI Prediabetes  UC Type 2 Diabetes | 230  96  66 | 1.00 (2.00)  1.00 (1.00)  1.00 (1.00) | 158  66  52 | 1.00 (2.00)  1.00 (1.00)  1.00 (1.50) | 72  30  14 | 1.00 (2.00)  1.50 (1.00)  1.00 (1.00) | 0.87  0.66  0.69 |
| **Sleep duration**  CCI Type 2 Diabetes  CCI Prediabetes  UC Type 2 Diabetes | 230  96  66 | 1.00 (2.00)  1.00 (2.00)  1.00 (2.00) | 158  66  52 | 1.00 (2.00)  1.00 (2.00)  1.00 (2.00) | 72  30  14 | 1.00 (1.00)  1.00 (2.00)  1.00 (1.00) | 0.95  0.60  0.46 |
| **Habitual sleep efficiency**  CCI Type 2 Diabetes  CCI Prediabetes  UC Type 2 Diabetes | 230  96  66 | 0.00 (1.00)  0.00 (1.00)  0.00 (1.00) | 158  66  52 | 0.00 (1.00)  0.00 (1.00)  0.00 (1.00) | 72  30  14 | 0.00 (1.00)  0.00 (1.00)  1.00 (1.00) | 0.48  0.66  0.09 |
| **Sleep disturbances**  CCI Type 2 Diabetes  CCI Prediabetes  UC Type 2 Diabetes | 230  96  66 | 2.00 (1.00)  2.00 (1.00)  2.00 (1.00) | 158  66  52 | 2.00 (1.00)  2.00 (1.00)  2.00 (1.00) | 72  30  14 | 2.00 (1.00)  2.00 (1.00)  2.00 (1.00) | 0.99  0.87  0.72 |
| **Use of sleep medication**  CCI Type 2 Diabetes  CCI Prediabetes  UC Type 2 Diabetes | 230  96  66 | 0.00 (1.00)  0.00 (1.00)  0.00 (2.00) | 158  66  52 | 0.00 (1.00)  0.00 (0.00)  0.00 (2.50) | 72  30  14 | 0.00 (1.50)  0.00 (2.00)  0.00 (0.00) | 0.27  0.07  0.34 |
| **Daytime dysfunction**  CCI Type 2 Diabetes  CCI Prediabetes  UC Type 2 Diabetes | 230  96  66 | 1.00 (1.00)  1.00 (1.00)  1.00 (1.00) | 158  66  52 | 1.00 (1.00)  1.00 (1.00)  1.00 (1.00) | 72  30  14 | 1.00 (1.00)  1.00 (0.00)  1.00 (1.00) | 0.56  0.57  0.64 |
